# Glycoinformatic profiling of label-free intact heparan sulfate oligosaccharides

**DOI:** 10.1101/2024.09.18.613784

**Authors:** Marissa L. Maciej-Hulme, Jandi Kim, Elijah T. Roberts, Yiqing Zhang, Anouk van der Velden, Dirk den Braanker, Cansu Yanginlar, Mark de Graaf, Ton Rabelink, Bernard van den Berg, Ellen van Omen, Rutger Maas, Anne-Els van de Logt, I. Jonathan Amster, Johan van der Vlag

**Author notes:** Contributed equally, in alphabetical order (Jandi Kim, Elijah T. Roberts, Yiqing Zhang).

## Abstract

Heparan sulfates (HS) are a group of heterogenous linear, sulfated polysaccharides that play a role in in health and many diseases including cancer, cardiovascular, and kidney diseases. The structural variety of HS has greatly challenged the development and utility of HS analytics, particularly for native structures, leaving a significant gap in HS technologies for clinical application. Mass spectrometry (MS)-based profiling with bioinformatics offers a top-down approach that can retain variety in large data sets. Using healthy human plasmas, we developed an MS glycoprofiling approach for native HS oligosaccharides, which retains the structural complexity of each individual HS chain and generates an HS ‘index’ (or Heparan-ome) for each patient. As a proof of concept, analysis of 56 plasma samples ranging from 6 groups of kidney disease patients revealed a new subset cluster (20%, 4/20) of membranous glomerulopathy (MG) patients with distinct HS profiles, highlighting the potential of HS glycoprofiling as a powerful new approach into clinical practice, which warrants future development into clinical diagnostics of kidney and other diseases.

**Graphical abstract:** 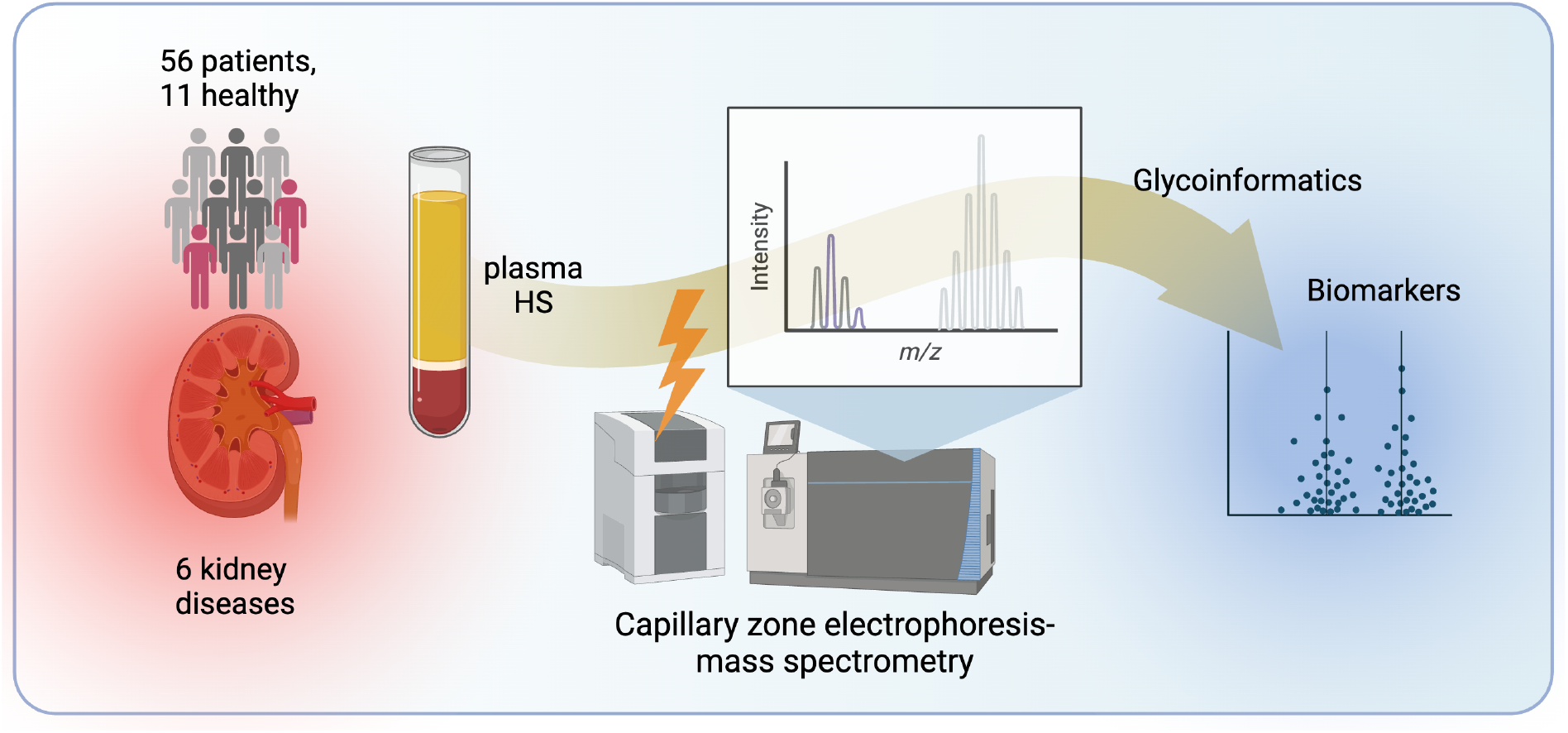

## Introduction

Glycans are a major class of molecules that are rich in information about the health status of an individual and play vital roles in protein interactions, stability, solubility, biological half life, and biological targeting, among others. Clinical interest in the glycome and glycomics has exponentially increased in recent years, owing to the rapid development of novel methods, tools, technologies and glycoinformatics. However, glycomics and glycoinformatics for certain types of glycans remain sparse. Glycosaminoglycans, such as heparan sulfates (HS), are particularly underrepresented in glycomics, hampered by inherent logistical challenges posed by their large size, structural similarities, high negative charge and labile sulfate groups. HS are a group of long, linear negatively charged polysaccharides decorated with sulfate groups and are found on the cell surface, in the extracellular matrix and in biofluids such as plasma and saliva. HS chains are synthesized in the Golgi of mammalian cells on serine residues of select proteins (termed proteoglycans, PGs). Each HS chain is built by a myriad of enzymes that collectively produce the characteristic heterogenic structures of HS. The positioning of sulfate groups along the HS chain give rise to distinct three-dimensional structural and chemical properties (Figure 1). Local biochemistry and architecture within the chain encode the interaction of HS with hundreds of different HS-binding proteins, enabling orchestration of a wide variety of biological activities including: 1) co-receptor roles in growth factor and cytokine signaling, 2) scaffolds for protein dimerization/multi-oligomerisation, and 3) in morphogen gradient dynamics and ligand sequestration mechanisms. HS heterogeneity is further expanded after biosynthesis at several points throughout its lifecycle, including potential modification by extra-Golgi sulfotransferases^1^, sulfatases and heparanase-1 (HPSE)^2^. Activated HPSE cleaves portions of HS chains; shortening the parent chain(s) left on the proteoglycan and releasing HS fragments into the extracellular milieu. Shedding of whole HSPGs from cells and the extracellular matrix can also occur during processes such as tissue remodeling, infection and inflammation^3^.

**Figure 1.**
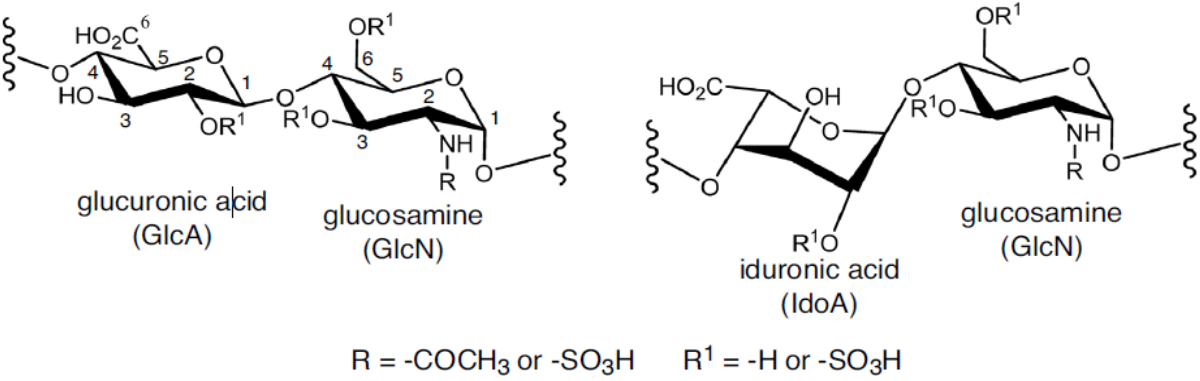
Heparan sulfate biosynthesis and structure.

Mass spectrometry (MS)-based glycomics offers an overview of structural glycan changes in a sample, providing potential insight into the considerable heterogeneity of HS by enabling multiple HS species from a single sample to be analysed simultaneously (profiling). Compositional analysis of plasma HS disaccharides by MS was recently demonstrated as a potential diagnostic for early stage cancer^4^, potentiating the use of HS glycoprofiling for personalized medicine from a simple blood draw. However, this method (and almost all clinical MS-based HS analyses conducted thus far) focused solely on the composition of HS disaccharides, which are derived from HS chains by enzymatic depolymerization using bacterial lyases. This step conveniently reduces sample heterogeneity to 12 distinct disaccharide species but simultaneously destroys vital information encoded within oligosaccharide architecture/sequence that influences HS biological function exerted through HS-binding partners. Since HS oligosaccharides contain more biological information than disaccharides, we sought to develop a novel and complete workflow pipeline from plasma sample preparation, through MS data acquisition, and onto glycoinfomatics for HS oligosaccharides. Our aim was to retain the natural heterogeneity of HS, as a more information-rich glycoprofiling approach for probing the health status of a patient, (*e*.*g*. a kidney patient). To demonstrate the concept, we purified and analysed plasma HS from healthy controls providing the first data set of plasma-derived biologically intact HS oligosaccharides.

We and others have previously established a prominent role for HS in the development of several kidney diseases^5-10^. In some of these, increased circulating plasma HS precedes and/or coincides with kidney damage, which suggests that plasma HS may be a potential biomarker for kidney injury and/or disease progression. Therefore, we collected plasma from 6 groups of kidney disease patients to see if analysis of HS oligosaccharides would identify disease-specific biomarkers. These data underpin the identification of plasma HS oligosaccharides associated with specific diseases and/or patients with potential disturbances in HS oligosaccharide structures.

## Results

First we collected plasma from healthy individuals and developed a sample preparation workflow compatible with either EDTA or citrate plasma (Figure 2). Sample preparation was adapted from previously established methods ^6,11,12^. Successful purification of plasma HS by the adapted method was confirmed by HS disaccharide analysis from 3 pooled healthy samples (Table 1). Total HS isolated from 11 healthy individuals varied from 2.25 µg/mL to 7.58 µg/mL, with an average of 5.37 µg/mL (Table 2). The number of detected species by capillary zone electrophoresis-mass spectrometry (CE-MS) did not correlate with the quantity of total HS (data not shown). Using in-house code, GAGWizard, information about each neutral mass was assigned including: mass to charge ratio (m/z), charge (z), intensity (I), migration time (MT) and neutral mass as well as the theoretical composition (degrees of polymerization (dp), number of hexaronic acids (#HexA), number of hexaronic amines (#HexN), number of sulfate groups (#SO3), number of acetyl groups (#NAc), the number and type of adduct, and elemental formula). HS dp size ranged between 2-34 and the number of sulfate groups varied from 0-29, demonstrating versatile detection of a structurally diverse range of HS species within each sample.

**Table 1.**
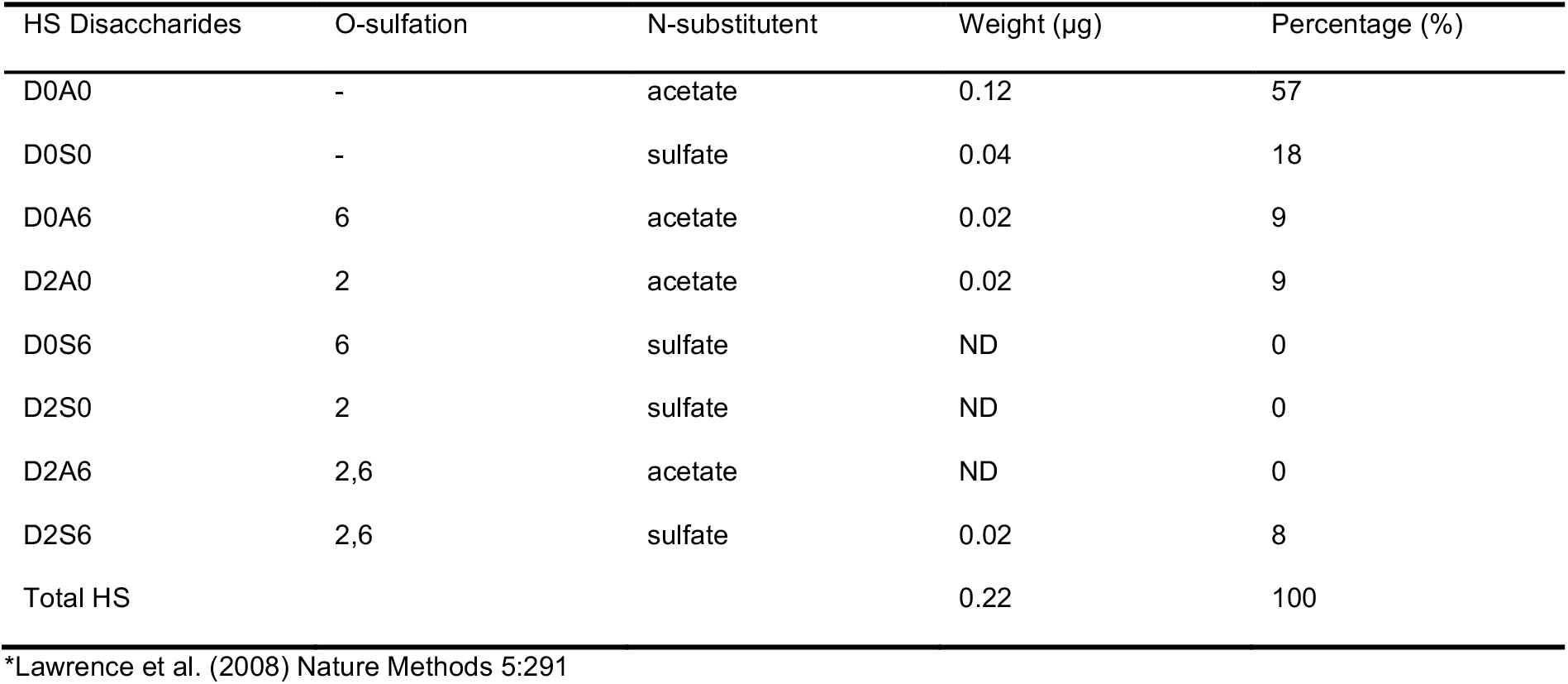
Heparin Disaccharide SAX-HPLC Analysis of Pooled Healthy Control Group.

| HS Disaccharides | O-sulfation | N-substituent | Weight (µg) | Percentage (%) |
| --- | --- | --- | --- | --- |
| D0A0 | - | acetate | 0.12 | 57 |
| D0S0 | - | sulfate | 0.04 | 18 |
| D0A6 | 6 | acetate | 0.02 | 9 |
| D2A0 | 2 | acetate | 0.02 | 9 |
| D0S6 | 6 | sulfate | ND | 0 |
| D2S0 | 2 | sulfate | ND | 0 |
| D2A6 | 2,6 | acetate | ND | 0 |
| D2S6 | 2,6 | sulfate | 0.02 | 8 |
| Total HS |  |  | 0.22 | 100 |

\*Lawrence et al. (2008) Nature Methods 5:291

**Table 2.** Healthy and patient metadata characteristics. *not detected, n.d. Kidney disease were grouped into proteinuric type 2 diabetes mellitus (T2D), membranous glomeruopathy (MG), systemic lupus erythematosus (SLE), focal segmental glomerulosclerosis (FSGS)/ minimal change disease (MCD) groups.

| Cohort | Number of participants | Mean µg/mL total HS (SEM) | HS dp size | Sulfates | Mean ImU/mL HPSE (SEM) |
| --- | --- | --- | --- | --- | --- |
| Healthy | 11 | 5.11 (1.75) | 2-34 | 0-29 | 0.13 (0.008) |
| FSGS,MCD | 15,2 | 9.77 (3.43)<br>*p*= 0.0002 | 3-22 | 0-29 | 0.22 (0.08)<br>*p*= 0.0074 |
| SLE | 11 | 3.61 (1.08) | 3-33 | 0-27 | 0.15 (0.034) |
| MG | 20 | 6.9 (2.55) | 2-33 | 0-29 | 0.13 (0.003)<br>*p* <0.0001 |
| T2D | 5 | nt | 2-9 | 2-11 | nt |

nt. not tested; n.s, not significant

**Figure 2.**
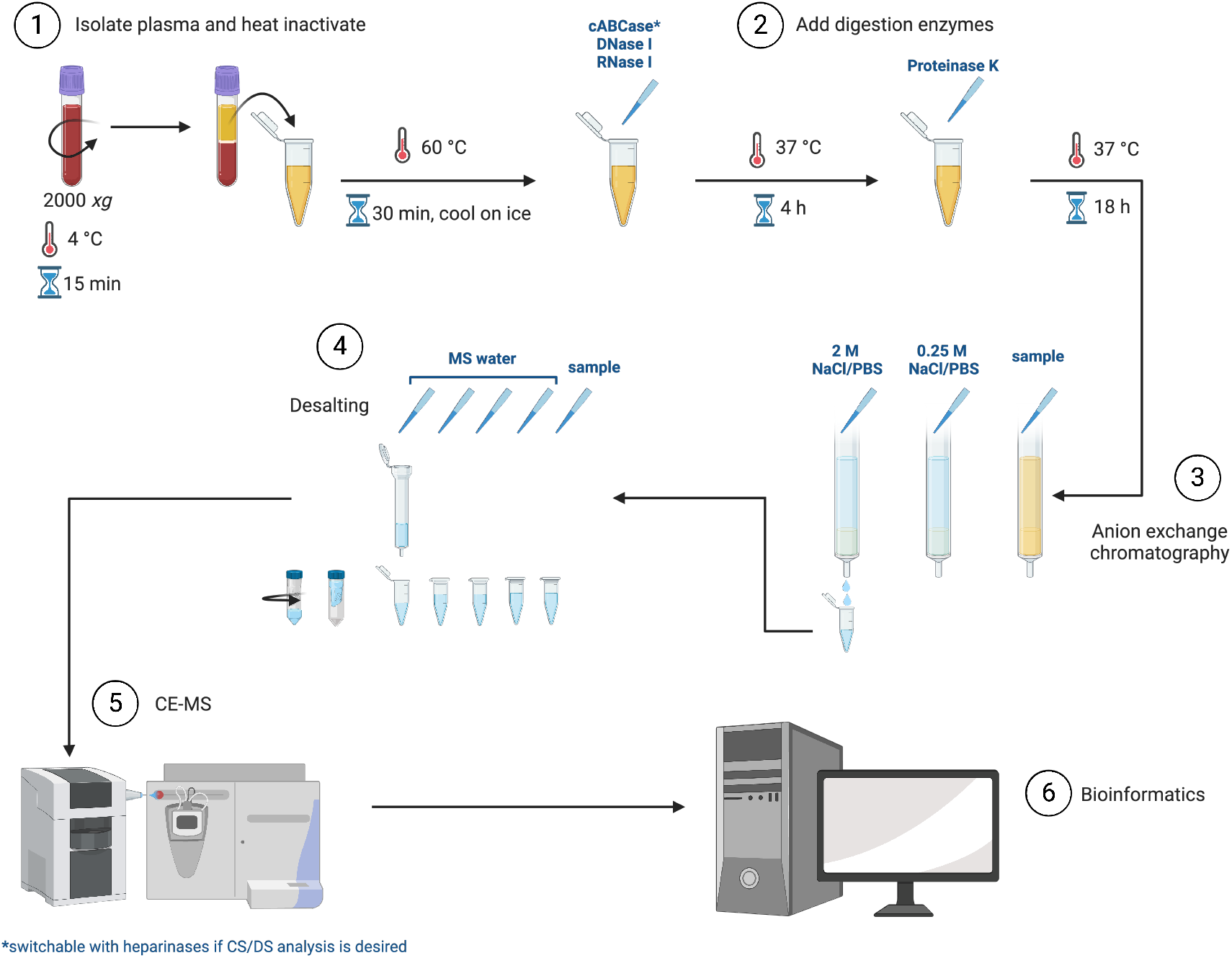
Plasma sample preparation overview for mass spectrometry compatible heparan sulfate isolation.

We then applied the same plasma purification and analysis method to 56 kidney disease patients diagnosed with either: proteinuric type 2 diabetes mellitis (T2D), membranous glomeruopathy (MG), systemic lupus erythematosus (SLE), focal segmental glomerulosclerosis (FSGS) or minimal change disease (MCD). Total HS quantities isolated from each kidney disease group were similar to healthy controls except for FSGS/MCD patients, which had significantly increased HS levels (Table 2). The size and total number of sulfates detected for almost all of the kidney diease groups were within the same range as HS extracted from healthy individuals. Only the T2D group had shorter oligosaccharides (dp 2-9) and less sulfates detected (2-11), whilst FSGS/MCD had slightly shorter HS oligosaccharides (dp 3-22).

Previously we successfully demonstrated HS oligosaccharide analysis on small scale sample sets from human urine^11,13^ and mouse cell culture preparations^6^ using a manual comparative approach. However application to larger datasets posed significant data processing problems due to a lack of appropriately tailored bioinformatic tools. Therefore we designed a data processing and glycoinformatics workflow for curated HS glycoprofiling suitable for large scale datasets (Figure 3). Once the matched HS lists for each plasma sample were obtained (steps 1-2 in Figure 3), data were further processed and curated to investigate the presence of any potential unique HS species within each patient compared with healthy samples. To do this, neutral masses from healthy plasma MS spectra were combined onto a master list (step 3), which was subsequently used in the data processing pipeline (step 4) to filter out healthy HS matches from kidney disease patient data. In step 5, data from the 3 technical replicate runs for each kidney disease patient plasma were condensed into one list per patient where each neutral mass species retained on the list appeared at least twice or more in MS spectra from replicate runs. Assignment of GAG compositions and further analysis were then based on these curated data (step 6-7).

**Figure 3.**
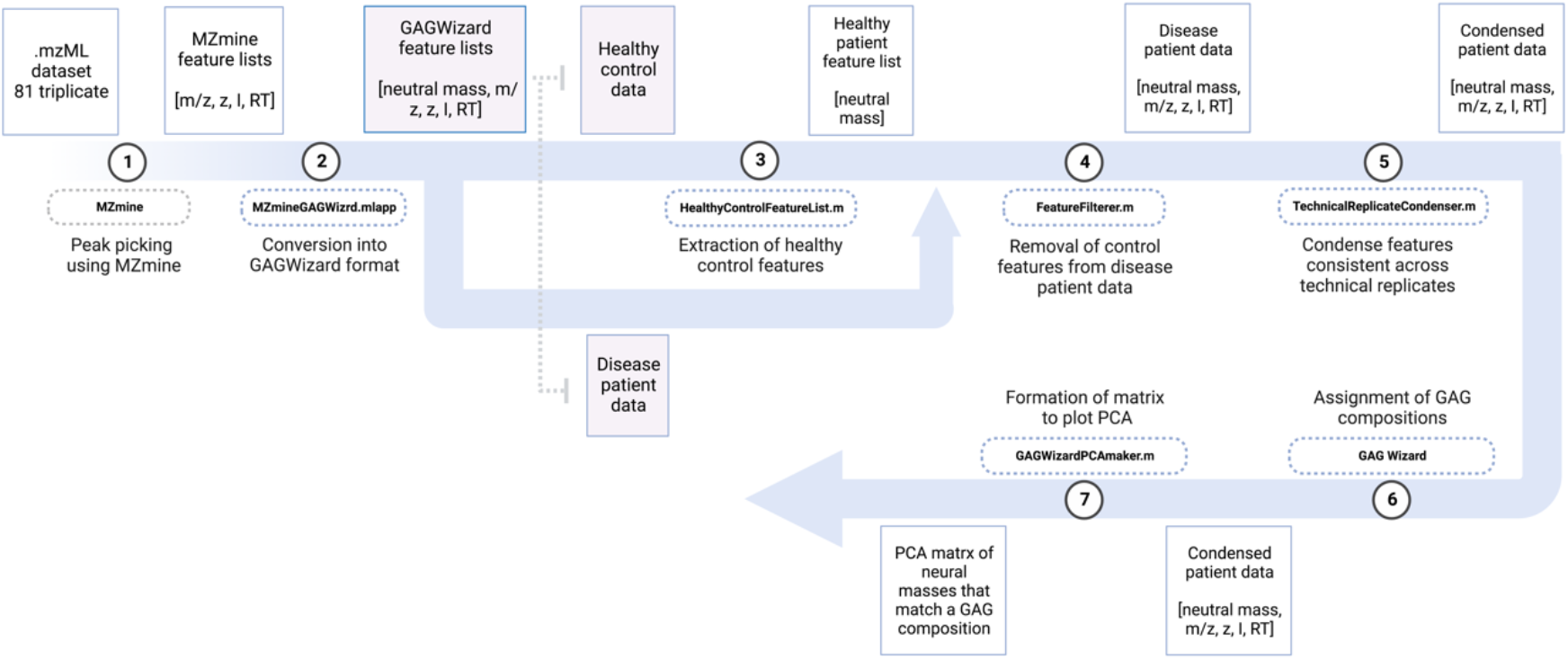
Data processing and bioinformatics workflow. Square boxes show collected data and their format styles. Dotted boxes with numbers inside of the blue arrow mean MATLAB scripts and the objectives. Steps 1∼2: MZmine and a MATLAB script provide raw data processing and GAGWizard feature lists. Steps 3∼4: Screening features that are only useful to the disease group using the healthy congrol filter script.Steps 5∼6: Completion of representative features, GAG composition assignment, and PCA application.

Grouped data for each disease showed shared and distinct features for each disease type; the most striking being for membranous glomerulopathy (MG), which contained the highest total number of HS species detected (88 species) of which 59 were unique to MG (Figure 4A). Each kidney disease group had at least 6 or more unique HS species. Multiple HS species were found in more than one disease group, with 2 HS species common to all kidney diseases (Figure 4B). Principle component analysis (PCA) analysis revealed a separating group from the majority of data where 4 MG patient HS profiles were clustered away from all other patients (Figure 4C). Upon closer analysis, these 4 patients had several similar sized HS oligosaccharides predicted and all shared 1 particular predicted neutral mass of 4343.511 (Figure 4D), with a suggested HS oligosaccharide structure depicted in Figure 4E. Multiple variations of the HS structure are possible but due to a lack of patient material, we were unable to assign the precise location of sulfate groups.

**Figure 4.**
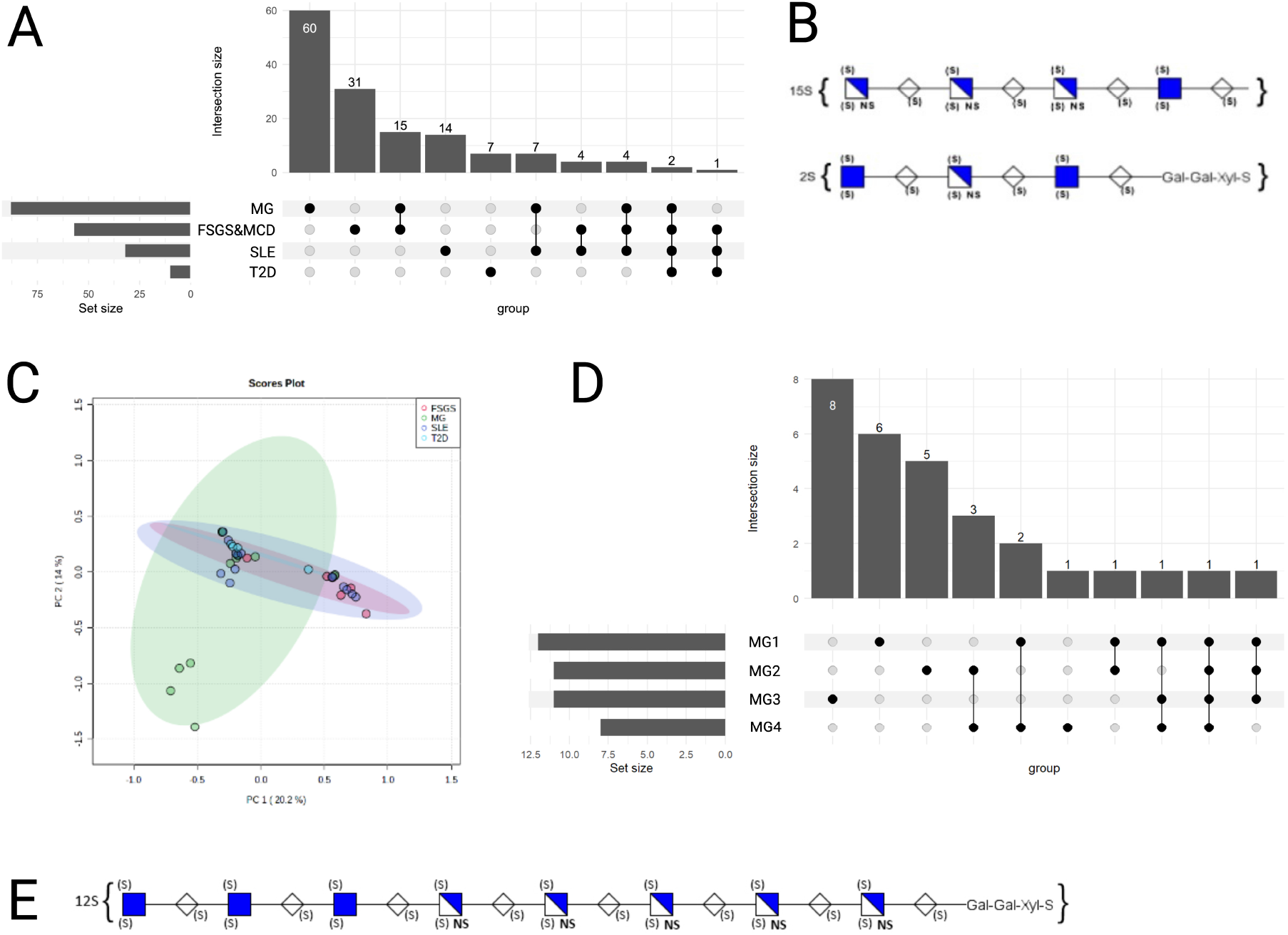
**(A) Upset plot of shared and unique HS hits for kidney diseases groups**. Bar chart represents number of HS species and intersections (connections) below bar chart shows connections of shared HS species between groups. **(B) Predicted structure of HS oligosaccharides shared between all kidney disease groups for each disease**. 2 neutral masses (1816.428 and 2607.777) common to all kidney diseases tested but not in healthy plasma were matched with predicted HS oligosaccharide structures. **(C) PCA of HS profiles for each disease**. PCA stratified data classifications based on disease type. Each dot represents a patient HS profile, with different colours representing different disease groups. Four MG samples were separated from other patients in which most of the variables were gathered in the PCA plot. **(D) Upset plot of HS oligosaccharides of MG subgroup identified by PCA in panel C**. 1 neutral mass (4343.511) was common to every MG subgroup. **(E) Predicted structure of common HS oligosaccharide in the MG sub group identified by PCA**.

One common mechanism that may modify the HS profiles of kidney disease patients is HPSE activity. Active HPSE degrades HS chains and releases HS oligosaccharide fragments from proteoglycans, and HS glycocalyx damage by HPSE plays a prominent role in the development of kidney disease^14^. Therefore, we also tested the plasma samples for HPSE activity to see if significant HPSE activity might produce a common HS oligosaccharide profile. Total HS quantification and HPSE activity data was not tested for T2D patients due to the lack of available plasma. Of the 3 other disease groups, only FSGS/MCD patients showed a significant increase in plasma HPSE activity accompanied by a significant increase in circulating total plasma HS (Table 2), and the presence of slightly shorter HS oligosaccharides (dp 3-22). In contrast, MG patients had statistically significantly lower plasma HPSE activity than healthy controls and total HS levels within the range of healthy controls. To investigate which HS species may be common to patients with increased HPSE activity, we excluded the T2D group and then stratified the patients irrespective of disease type into 2 groups: HPSE_nor(mal)_ and HPSE_high_ groups, where HPSE_high_ represented patients with enzyme activity values above those found in healthy controls (Figure 5). 21 HS species were found only in HPSE_high_ patients, whilst 20 were shared between HPSE_high_ and HPSE_nor_. The largest number of species was associated with HPSE_nor_ alone (104 species), suggesting that the majority of plasma HS oligosaccharides are not associated with HPSE activity. No distinctive separation occurred following PCA analysis (data not shown) between HPSE_high_ and HPSE_nor_ groups.

**Figure 5.**
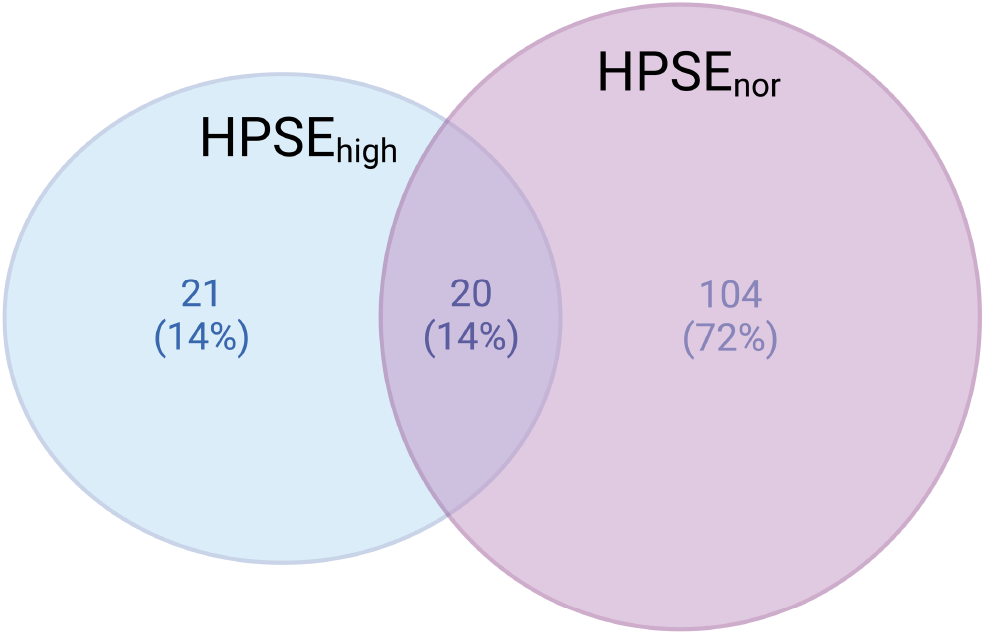
Venn diagram of shared and unique HS hits for HPSE_nor_ and HPSE_high_ groups.

## Discussion

Kidney disease affects more than 10% of the global population (>800 million individuals) and represents one of the leading causes of mortality worldwide, with associated deaths continuing to rise. Although biopsy and subsequent histological analyses remain the gold standard for kidney disease diagnosis, the discovery of biomarkers derived from less invasive techniques, such as biofluids, would increase comfort for the patient, reduce the burden on the healthcare labour force thereby reducing costs and potentially highlight underlying mechanisms for novel treatment strategies. Human plasma is a biofluid rich in information about patient health status and a routinely biobanked sample. Plasma offers a liquid biopsy option of diverse sample collections from a plethora of diseases and easy integration into diagnostic processes in healthcare systems. Various diseases have been reported to show increases in circulating plasma HS, including COVID-19^15^, sepsis^16^, cancer^4^, and kidney diseases (Table 2,^17,18^). Current heparan sulfate purification methods for MS glycomics rely on simplifying the complexity of heparan sulfates to determine disaccharide composition, which does not identify the actual sequence information that underpin biological function. Here we took a different approach and developed an MS-compatible preparation method and systematic glycoinformatics of biologically relevant HS oligosaccharides resulting in individual glyco-’fingerprints’ for patients. This method represents a core method for application to almost any plasma sample (excluding heparinised plasma due to the presence of heparin) derived from patients with diverse conditions or diseases. The method can easily be adapted to profile chondroitin/dermatan sulfate through omission of the cABCase buffer and exchange of chondroitinase ABC for heparinase enzymes (I-III)^12^, extending the utility of the approach.

As a proof of concept for our novel methodology, we analysed plasma HS from healthy controls and a range of kidney diseases to probe the utility of the method for disease stratification and to develop a bioinformatics workflow compatible with clinical demand for larger datasets. Using this approach, we discovered several unique and shared HS species among the different kidney diseases in our analyses. Shared HS species between kidney disease groups may be indicative of general mechanisms such as acute inflammation that result in tissue injury when not resolved, whereas unique HS species may represent potential biomarkers for specific kidney disease diagnostics. Specifically, our approach identified 2 common HS oligosaccharides for kidney disease that were not found in healthy plasma, representing two potential biomarker candidates for general kidney disease detection via a non-invasive diagnostics approach from patient plasma. In addition, our methodology also identified a unique subset of patients with MG, which is an autoimmune kidney disease characterized by proteinuria, deposition of immune complexes in the kidney and thickening of the glomerular basement membrane. Interestingly, recent data has pointed towards a role for HS in disease development and/or progression of MG. Antibodies in MG patients to HS have been found^19^, as well as the abnormal presence of the HS copolymerase complex enzymes, EXT1 and EXT2, in the glomerular basement membrane^20^. Furthemore, two of the established autoantibody targets for MG, Phospholipase A2 receptor and (PLA2R) and thrombospondin type 1 domain containing 7A (THSD7A), both have associations with HS. PLA2R interacts with the established HS binding protein, PLA2^19^, and THSD7A is predicted to have an HS binding site^21^. Our data suggests that HS oligosaccharides themselves may be dysregulated, producing distinct profiles for a subset of MG patients. Together, these phenomena suggest that disturbances in HS expression, modification and localisation may underpin facets of MG.

In summary, we describe a cutting-edge method and data processing pipeline for the profiling of intact HS oligosaccharides from the plasma of healthy and kidney disease patients. Using these samples, we demonstrate how plasma HS glycoprofiling by MS and glycobioinformatics generates an individual glyco-’fingerprint’ for patients that may serve, along with other parameters, as a future clinical diagnostic tool. Our analyses also highlight particular HS species as potential non-invasive novel biomarkers (liquid biopsy) for general kidney disease detection and in a subset MG patients that warrant further exploration.

## Methods

### Preparation of human plasma and HS oligosaccharide purification

Written informed consent was obtained from healthy volunteers and patients prior to blood collection. EDTA-collected blood was centrifuged at 4^°^C for 15 mins at 2000 *xg*. The plasma (supernatant) was heat inactivated at 60^°^C for 30 mins, then rapidly cooled on ice. 1 mL plasma was supplemented with cABCase buffer (25 mM Tris, 2 mM Mg(Ac)_2_ pH 8), then treated with 2 mU of chondroitinase ABC (Sigma), 7.5 U/mL DNAse I (Qiagen) and 10 U/mL RNAse I (ThermoScientific) for 4 hours at 37^°^C, followed by addition of 125 µg/mL proteinase K (Merck chemicals B.V., Amsterdam, The Netherlands) and continued digestion at 37^°^C overnight. To inactivate proteinase K, samples were heated at 95^°^C for 10 mins, then cooled on ice. Samples were 5x diluted with room temperature (RT) sterile PBS immediately prior to HS isolation to adjust the temperature and pH. Anion exchange chromatography was performed using individual 1 mL bed volume DEAE-sepharose CL-6B beads (Sigma) prepared as previously described^12^ and equilibrated in PBS at RT. Bound HS was washed with 0.25 M NaCl/PBS, pH 7.4 and then eluted with 4x 1 mL of 2 M NaCl/PBS, pH 7.4. Isolated HS was desalted in 1 mL aliquots using PD10 desalting columns (GE Healthcare, Sephadex G25) with CHROMASOLV™ LC-MS H_2_O (Fisher Scientific), dried by centrifugal evaporation and stored at -20^°^C.

### HS quantification

Isolated HS was quantified in black walled 96-well microplates for fluorescence-based assays, (Invitrogen) using the HeparinRed assay (Red Probes, Germany) according to the manufacturer’s instructions. Porcine HSBK (Sigma) was used to prepare the standard curve. Fluorescence was measured after linear shaking for 20 secs (_ex_ 570 nm, _em_ 605 nm) using an Infinite M200 Pro fluorescence plate reader and software.

### HPSE activity assay

Plasma HPSE activity was measured using the Heparan degrading enzyme assay kit (Takara Bio, #MK412) following the manufacturer’s instructions.

### CE-Orbitrap MS analysis

Capillary zone electrophoresis (CE) (Agilent G1600 HP 3D CE) was coupled to an Orbitrap Elite mass spectrometer (Thermo Fisher Scientific, Bremen, Germant) with an EMASS-II CE-MS Ion Source (CMP Scientific, Brooklyn, NY). Mass spectra were collected in negative mode at a resolution of 120,000 at m/z 200, with one micro scan with a maximum injection 200 ms. All mass spectra were collected in profile mode. The automatic gain control (AGC) target value was set to 1e^6^. Prior to CE-MS experiments, a semi-automatic optimization of source parameters was performed using sucrose octasulfate to improve sensitivity of sulfated GAGs and reduce sulfate decomposition during ion transfer prior to MS analysis. To minimize sulfate loss, the source region stacked-ring ion guide (S-lens) was set to 20%. The ion transfer tube was maintained at ground potential and at 200^°^C. CE separations were performed on fused silica capillaries (60 cm x 360 µm OD x 50 µm ID) functionalized with dichlorodimethylsilane (DMS) using 25 mM ammonium acetate in 70:30 MeOH:H_2_O as a background electrolyte. DMS functionalization and HF etching procedures are reported previously^11,22^. The outlet end of the capillaries were etched with hydrofluoric acid (HF) to reduce the outer diameter to <100 µm and improve performance of the EMASS-II sheath flow interface. Glass electrospray emitters with ∼30 µm orifice were used in the interface, which were filled with 25 mM ammonium acetate in 70% (v/v) methanol/water which acted as a sheath liquid ^23^. The glass emitter was fabricated with Sutter P-1000 pipette puller (Sutter instrument, Novota, California) in the UGA Biomembranes Engineering Facility. The etched capillary was positioned 0.3-0.5 mm from the end of the emitter orifice to create a mixing volume of approximately 2 nL. The small mixing volume significantly reduces the dilution of analytes, compared to the conventional CE-MS sheath interface. The glass emitter was aligned ca. 2.5 mm away from the inlet capillary of the orbitrap. A voltage of - 1.70 to -1.85 kV was applied to the sheath liquid (SL) to achieve electrospray.

Dry HS samples were dissolved in 400 µL HPLC grade water and further de-salted with 0.5 mL 3 K Amicon Ultra spin filters. The cellulose membrane filter was washed with H_2_O 1 min before application of the rehydrated samples, which were centrifuged three times at 14,000 *xg* for 25 mins. Each desalted aqueous GAG sample was transferred to a 250 µL CE vial with a pre-slit septum cap on. The aqueous GAG samples were injected for 6 s at 950 mbar followed by a background electrolyte (BGE) injection for 10 s at 50 mbarfor a total injection volume of 156 nL (sample plug was 14.5% for a 60 cm length column). The HS samples were run in triplicate in a randomized sequence to minimize system bias^24^. Every tenth sample injection was followed by a blank injection and standard Dermatan Sulfate to monitor the system for carryover and mass accuracy shifts. Total CE-MS running time was set to 60 min. After each run, 5 min fresh BGE flush was followed to remove residual contaminants in the capillary.

### Data processing of Orbitrap mass spectra

Mass spectra were visualized in Thermo Xcalibur 2.2 software. MS convert (developed by Jerry Holman^25^) in ProteoWizard 3.02 package was utilized to convert Thermo raw files to open-format mzML files. None of the filters are applied in MS convert GUI. Binary encoding precision was 32 bits. Zlib compression was used for file size deduction and TPP compatibility was allowed.

MZmine 3.1 package was employed to extract MS features (m/z, charge, retention time, and intensity)^26^. Part of key parameters used in MZmine 3.1 were optimized with Paramounter R package developed by Guo^9^. The local minimum algorithm was selected for the deconvolution. The setting of MZmine 3.1 parameters is described below. Mass tolerance for chromatogram builder was assigned to ±5 parts per million mass resolution to seek an accurate mass m/z. The wavelet transform was selected as the mass detector with scale level 5 and wavelet window size 30%. The signal/noise threshold was set to the estimation of local minima by in-house software^27^. Next, the ADAP chromatogram builder parameters were set as follows: minimum absolute peak height 2.0 × 10^2^ and minimum number of scans 5. Electropherogram deconvolution parameters were established as follow: Chromatographic threshold 85%, minimum peak top/edge ratio 1.7, minimum absolute intensity 4.0 × 10^2^, and maximum peak duration range 10 min. Isotopic peaks grouper algorithm was chosen to deisotope features with retention time tolerance 0.41 min, maximum charge five, and the lowest isotope was selected, which is the monoisotopic peak.

### Healthy control filter

In house software (GAGWizard) was used to identify features that were associated with healthy control patients. To meet this criteria, a neutral mass feature must appear in at least two technical replicates of a healthy patient, and only has to appear in at least one patient sample. These features were then removed from the remaining datasets for patients.

### Technical Replicate Condenser

For each set of technical replicates (in triplicate), the feature lists were condensed into a single list. Features that were found in at least two of the three technical replicates were allowed to survive on the condensed list, and a 5 ppm error tolerance was used for neutral mass variation. The neutral masses and intensities were saved on the final list.

### Common data processing; MZmine2GAGWizard

MZmine2GAGWizard is an in house MATLAB script that calculates the neutral mass for each feature based on its m/z and charge, and also reorders the columns [neutral mass, m/z, charge, intensity, MT]. Neutral mass = (m/z * charge) + (1.0078*charge)

### GAGWizard: HS oligosaccharide composition predictions

Likely HS type GAG compositions were found for the remaining neutral mass features using GAGWizard. GAGWizard generates a combinatorial database of theoretical GAG compositions and their neutral masses which can be used to search against the data. For this study, a database of HS type GAGs was generated to resemble fully intact HS that are found on HSPGs. Compositions were generated with degrees of polymerization between dp2 and dp50. “Gal-Gal-Xyl-Ser” was tenable linker region considered because of the proteinase K digestion in this protocol. This generated a list of 35,444 theoretical compositions ranging from 940 Da to 16.9 kDa.

The mass lists for each patient were uploaded to GAGWizard, and the software searched the database for theoretical masses that matched the data within 5 ppm mass error. If multiple compositions fell within the 5 ppm error tolerance, each of them was returned on the results file. The output of GAGWizard contains information about the input data (m/z, z, I, MT, neutral mass) as well as information about the theoretical composition that it matched, which includes DP, #HexA, #HexN, #SO_3_, #NAc, the number and type of adduct, as well as the elemental formula. Lastly the ppm error is calculated for each matching composition.

### Principal Component Analysis (PCA)

Neutral masses that were found to match known GAG compositions were extracted from the GAGWizard results files and collated into a list. A PCA matrix was constructed with MATLAB where each row is a neutral mass feature, and each column represents a patient. The intensity of every neutral mass feature was filled in the middle of the matrix. If a neutral mass feature was not found for a particular patient, that cell was filled with a zero, and if a neutral mass matched multiple HS compositions, that neutral mass was only added to the matrix once. The matrix was uploaded to MetaboAnalyst which is a web-based platform for comprehensive data analysis^28^. Pre-processing steps included mean centering and normalization by the highest peak intensity for each neutral mass. Considering that there are a significant number of variables in Orbitrap dataset, inbuilt interquartile range (IQR) filter of Metaboanalyst was automatically applied to keep only top maximum features, which is strongly recommended for untargeted metabolomics dataset.

### HS biomarker discovery

Neutral mass lists with GAG composition matches grouped by disease were collated into one neutral mass list per disease group. To make the HS hits database, collated neutral mass lists were input into Venneuler R package and hits were calculated from the intersections of the neutral mass list. The UpSet diagram represents unique and shared HS hits within kidney disease cohorts, sorted by frequency-proportional manner.

## Data availability

Raw data files are available on request.

## Code availability

Custom computer codes used in this work are available on request.

## Conflict of Interest

The authors declare that the research was conducted in the absence of any commercial or financial relationships that could be construed as a potential conflict of interest.

## Acknowledgements

This study was supported by the consortium grant LSHM16058-SGF (GLYCOTREAT; a collaboration project financed by the PPP Allowance made available by Top Sector Life Sciences & Health to the Dutch Kidney Foundation to stimulate public-private partnerships) and a Marie Curie Fellowship GlycoMap #101107665 awarded to MLMH. Figures 2 and 3 were created with BioRender.

## Author contributions

MLMH conceived the hypotheses, designed, and conducted experiments, collected plasma samples, analysed data and co-wrote the first draft of the manuscript. JK performed experiments, analysed data and co-wrote the first draft of the manuscript. YQ performed experiments and analyzed data. ETR analyzed data. IJA and JV designed experiments and analyzed data. AV, DB, CY, MG, TR, BB, EO, RM, and AEL collected and curated plasma samples, and analyzed data. All authors contributed to manuscript revision, read and approved the submitted version.

## Notes

### Competing Interest Statement

The authors have declared no competing interest.

